# Low gH/gL (sub)species-specific antibody levels indicate elephants at risk of fatal elephant endotheliotropic herpesvirus hemorrhagic disease

**DOI:** 10.1101/2023.06.15.545196

**Authors:** Tabitha E. Hoornweg, Willem Schaftenaar, Victor P.M.G. Rutten, Cornelis A.M. de Haan

**Affiliations:** Section of Virology, Division Infectious Diseases and Immunology, Faculty of Veterinary Medicine, Department of Biomolecular Health Sciences, Utrecht University, Utrecht, The Netherlands; Section of Immunology, Division Infectious Diseases and Immunology, Faculty of Veterinary Medicine, Department of Biomolecular Health Sciences, Utrecht University, Utrecht, The Netherlands; Veterinary Advisor EAZA Elephant TAG, Rotterdam Zoo, Rotterdam, The Netherlands; Faculty of Veterinary Science, Department of Veterinary Tropical Diseases, University of Pretoria, Onderstepoort, South Africa

**Author notes:** Address correspondence to Tabitha E. Hoornweg, and C.A.M. (Xander) de Haan.

## Abstract

Elephant endotheliotropic herpesviruses (EEHVs), of which seven species and multiple subspecies are currently distinguished, naturally infect either Asian (*Elephas maximus*) or African elephants (*Loxodonta* species). While all adult elephants are latently infected with at least one EEHV (sub)species, EEHV infections in young elephants may lead to acute, fatal hemorrhagic disease (EEHV-HD). The disease is primarily observed in calves with low to non-detectable EEHV-specific antibodies, suggesting it is the result of an uncontrolled primary infection. Yet, young elephants showing high antibody levels in serological assays based on EEHV(1A) gB, which detect antibodies against multiple EEHV (sub)species, are not necessarily protected against EEHV-HD. To better determine which animals are at risk of EEHV-HD, we developed gB and gH/gL ELISAs for all Asian elephant EEHV (sub)species and assessed their specificity using 396 sera from 164 Asian elephants kept in Europeans zoos. Antibody levels measured against gB of different EEHV (sub)species correlated strongly with one another, indicating gB-specific antibodies are highly cross-reactive. In contrast, antibody responses towards gH/gL of different EEHV (sub)species were far less correlated and could be used to distinguish between infections with different (sub)species. Subsequently, antibody responses in sera of 23 EEHV-HD fatalities were analyzed. Whereas high antibody levels against gB could be detected in sera of multiple EEHV-HD fatalities, all fatalities had low antibody levels against gH/gL of the EEHV (sub)species they succumbed to. Overall, our data indicate that (sub)species-specific gH/gL ELISAs may be used to identify animals at risk of developing EEHV-HD upon infection with a particular EEHV (sub)species.

## Introduction

Elephant endotheliotropic herpesviruses (EEHVs) are elephant-specific viruses that may cause an acute, highly lethal disease in their natural host, known as EEHV-hemorrhagic disease (EEHV-HD). Currently seven EEHV species (EEHV1-7) are distinguished, of which EEHV1, -3, -5 and -7 are further subdivided into (sub)species A and B. Whereas EEHV species 1, 4, and 5 naturally infect Asian elephants (*Elephas maximus*), the two species of African elephants (*Loxodonta africana* and *Loxodonta cyclotis*) are natural carriers of EEHV2, -3, -6, and -7 (1).

All adult elephants tested to date were observed to have antibodies to EEHVs (2–5), suggesting they have been infected with either one or more EEHV (sub)species during their lives. Analogous to other herpesvirus infections, EEHVs are considered to stay latently present in infected animals, being occasionally shed as part of a natural infection cycle, typically without clinical signs (6, 7). In contrast, young elephants, usually between one and ten years of age, may develop EEHV-HD in response to EEHV infection. The disease is highly lethal and progresses quickly; calves often die within 24 to 48 hours after onset of symptoms (1, 8). Over the last 35 years, 12-17% of all Asian elephant calves born in Western zoos succumbed to EEHV-HD before reaching adulthood (8–10), with EEHV1A responsible for the majority of the fatalities. EEHV-HD has shown to also affect both (semi-)captive and free-ranging Asian elephants in range countries (11–13). As yet the exact mortality rates in these populations are unknown. Although the disease was long thought to solely affect Asian elephants, recently multiple young African elephants born in Western zoos succumbed to EEHV-HD as well (4, 14).

Several studies showed that animals with low to non-detectable EEHV-specific antibody levels are the ones at risk of EEHV-HD (2–5), suggesting that EEHV-HD may be the result of an insufficiently controlled primary EEHV infection. As calves below one year of age, which have comparable high EEHV-specific antibody levels to their dam (2, 15), rarely develop EEHV-HD and since risk of EEHV-HD greatly increases upon waning of maternal immunity (5), maternal antibodies are now thought to play a critical role in protection against EEHV-HD. Even though based on a limited number of animals, previous studies (2, 5) suggest that a past infection with one EEHV (sub)species does not necessarily protect against EEHV-HD upon subsequent infection with another EEHV (sub)species. In order to address this notion and to determine which animals are at risk for developing EEHV-HD, serological assays are needed that are able to differentiate between infection(s) with the different EEHV (sub)species.

For EEHV1A and EEHV1B, two of the EEHV subspecies infecting Asian elephants, differentiating serological assays have been developed employing the EEHV1-specific ORF-Q protein (2). However, a recent study described that some infected animals do not seroconvert to the ORF-Q antigen used in these serological assays (16). More importantly, differentiating serological assays are still lacking for the other EEHV (sub)species infecting Asian elephants (EEHV4 and -5). We previously described development of EEHV-specific ELISAs using EEHV1A glycoprotein B (gB) and the glycoprotein H/glycoprotein L heterodimer (gH/gL) as antigens, and identified gH/gL to be a promising candidate antigen for development of ELISAs that can differentiate between the different EEHVs (sub)species (3). In the present study, we developed gB and gH/gL ELISAs for all Asian elephant EEHV (sub)species and tested their reactivity and specificity using a large panel of elephant sera. While high antibody levels against gB were not necessarily indicative of protection against EEHV-HD, fatal EEHV-HD cases never had high antibody levels against gH/gL of the EEHV (sub)species they succumbed to. These results indicate that our gH/gL ELISAs are able to identify animals at risk of developing EEHV-HD, even if these have been infected with one or more EEHV (sub)species before.

## Materials and Methods

### Serum samples

A total of 396 serum samples from 164 individual Asian elephants (*Elephas maximus*), aged 0 days to 57 years, from 29 European zoological collections were used in this study. All blood samples were taken aseptically from ear or leg veins by zoo veterinary staff (as part of routine management recommended by the elephant Taxon Advisor Group of the European Association of Zoos and Aquaria) and sera were transported at 4°C to our institute for diagnostic purposes. Sera were stored at −20°C until use. For a number of analyses, subsets of the available serum samples were used as indicated in the respective figure legends.

### Phylogenetic analyses

Full length gB, gH and gL amino acid sequences of each EEHV (sub)species were retrieved from GenBank (February 22, 2023; n = 9 for gB and n= 8 for gH and gL, respectively). Sequences of gH and gL were concatenated per viral strain, and gB as well as gH/gL sequences were aligned using Clustal Omega. For both alignments, IQTree web server (17) was used to select the best-fit evolutionary model and infer a maximum likelihood (ML) tree. Robustness was assessed by ultrafast bootstrapping using 1000 replicates, and eventual ML trees were visualized and edited using FigTree (18). Identity matrices generated by clustal omega were edited in Microsoft Excel.

### Expression of recombinant EEHV proteins

Protein expression constructs for gB, gH and gL of EEHV (sub)species 1B, 4 and 5A were designed as previously described for EEHV1A (3). Details of the constructs are summarized in Table 1. Codon-optimized cDNAs (Genscript Biotech, Leiden, The Netherlands) were cloned into a pFRT expression plasmid (Thermo Fisher Scientific) in frame with an N-terminal Gaussia luciferase (Gluc) signal sequence and a C-terminal triple StrepTag (3xST; gB and gH constructs) or HisTag (6xHis; gL constructs) (3).

**Table 1.**
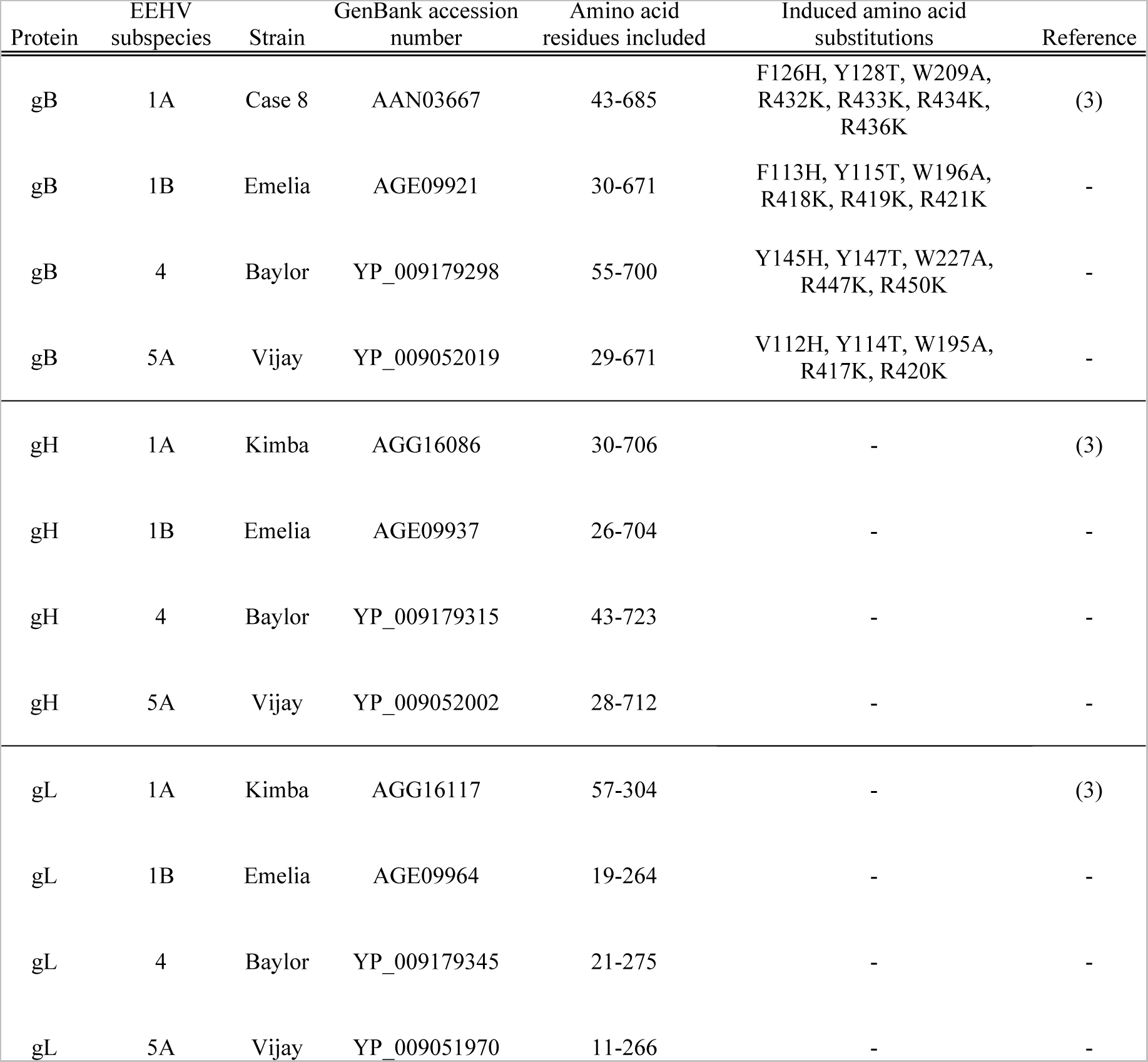
Protein expression constructs.

Individual plasmids or combinations of (sub)species-specific gH and gL expression plasmids were transfected into FreeStyle 293-F cells (Thermo Fisher Scientific) using polyethylenimine (PolySciences). Five days post transfection cell culture media were harvested, cleared of debris by low speed centrifugation after which proteins were purified using Strep-Tactin Sepharose beads (Iba). Purity and integrity of all proteins were checked and protein concentrations estimated using quantitative densitometry on GelCode Blue-stained protein gels (Thermo Fisher Scientific) containing bovine serum albumin (BSA) standards. Signals were imaged and analyzed using the Odyssey imaging system (LI-COR). When indicated, proteins were deglycosylated by PNGaseF (NEB) prior to gel electrophoresis to facilitate analysis.

For Western Blot analysis, proteins were transferred to a PVDF membrane using the Transblot Turbo System (BioRad) and subsequently stained using a horseradish peroxidase (HRP)-conjugated monoclonal anti-StrepTag antibody (Iba) or mouse anti-6His (Clontech) monoclonal antibody combined with rabbit anti-mouse-HRP polyclonal antibodies (DAKO). Signals were detected by use of enhanced chemiluminescence (ECL) Western Blotting substrate (Pierce, Thermo Fisher Scientific) and the Odyssey imaging system (LI-COR).

### ELISAs

All ELISAs were performed using protocols previously described for EEHV1A gB and gH/gL ELISAs (3). To ensure coating of equal amounts of gB or gH/gL proteins of different (sub)species, antigen dilution ranges were coated onto ELISA plates and the amounts of proteins needed to obtain equal OD values when probed using the HRP-conjugated monoclonal anti-StrepTag detection antibody (Iba), was determined. Coating ratios were determined using the following formula: (ng protein coated for EEHVx to obtain an OD of y)/(ng protein coated for EEHV1A to obtain an OD of y). Eventual coating concentrations for each antigen were calculated by multiplying its respective coating ratio with the optimal coating concentration previously established for EEHV1A gB (5 ng/well) respectively gH/gL (40 ng/well) (3). For all ELISAs, ΔOD values (OD value antigen coated well - OD value uncoated well) are reported as described previously (3). To facilitate comparison of ΔODs obtained in the different gB respectively gH/gL ELISAs, all ΔOD values were normalized relative to results obtained with a set of control sera.

### Statistical analyses

Mann-Whitney tests and pairwise simple linear regression analyses were performed using GraphPad Prism software. P values ≤0.05 were considered significant.

## Results

### High levels of EEHV gB-specific antibodies found in sera of several EEHV-HD fatalities

While elephants with low to non-detectable EEHV-specific antibody levels are known to be at risk for developing fatal EEHV-HD, it was previously noted that not all animals with high gB-specific antibody levels were protected against EEHV-HD (2, 5). To substantiate this finding, we compared gB-specific antibody levels of 72 Asian elephants below the age of 10, that either did not (yet) develop EEHV-HD (Figure 1A; n=49) or did develop fatal EEHV-HD (Figure 1B; n=23), plotted according to age at sampling. Even though antibody levels of EEHV-HD fatalities were significantly lower than those for non-HD elephants (Figure 1C), presence of high levels of EEHV-specific antibodies was not necessarily indicative of protection against EEHV-HD (Figure 1B). Notably, two EEHV-HD fatalities (Figure 1B; shown in red and light green) showed a clear increase of EEHV-specific antibody levels during their lives, strongly suggestive of a past EEHV infection, yet still succumbed to EEHV-HD.

**Figure 1.**
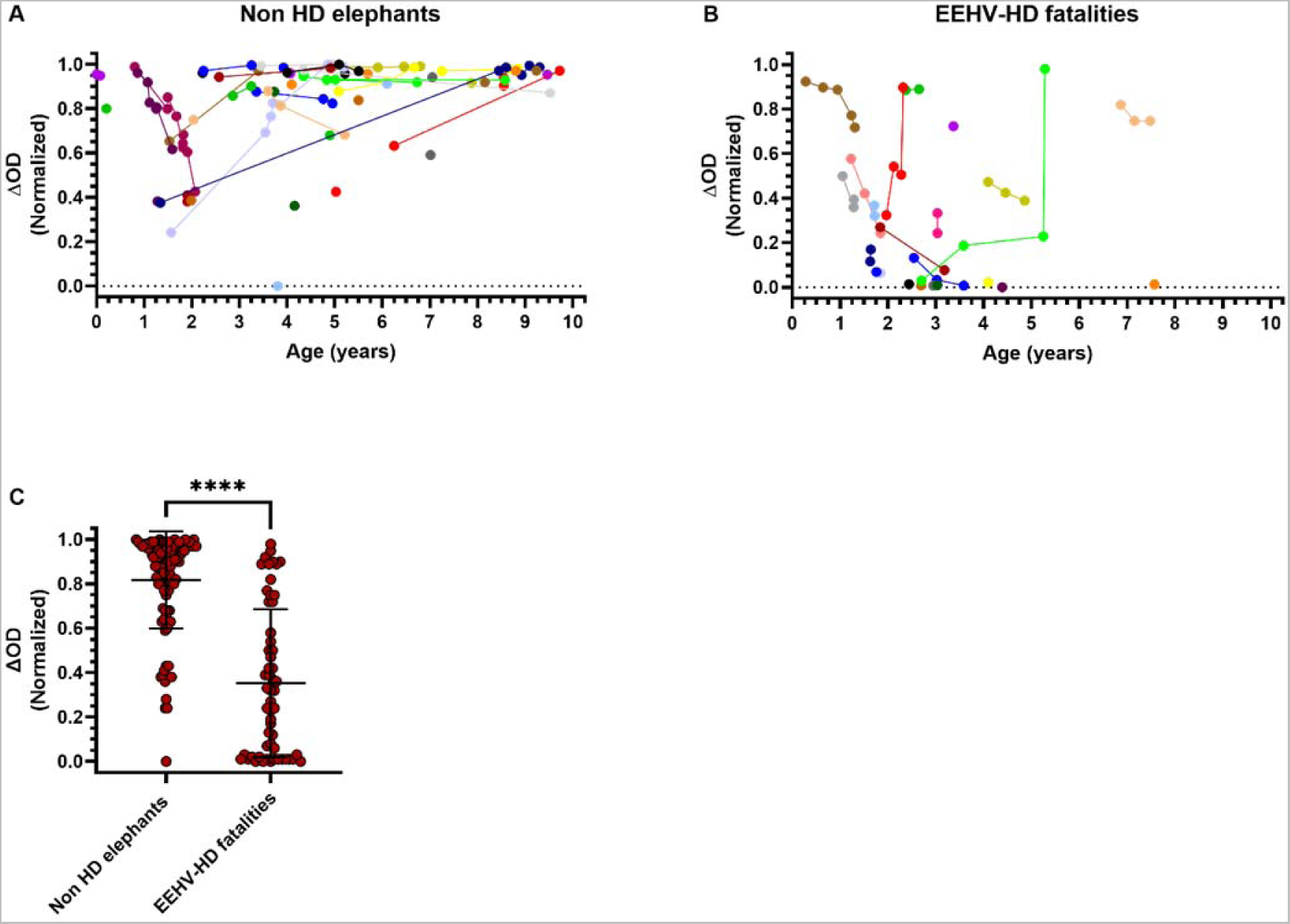
EEHV gB-specific antibody levels in young Asian elephants that did not develop EEHV-HD as compared to EEHV-HD fatalities. EEHV-specific antibody levels measured using the multiple EEHV species (EEHV1A) gB ELISA for a cross-sectional cohort of 102 serum samples from 49 individual Asian elephants below the age of 10 that did not (yet) develop EEHV-HD (A) and 56 serum samples from 23 individual EEHV-HD fatalities (B). All samples were tested at 1:100 dilution and ΔOD values were obtained by subtraction of serum-specific background signals from gB-specific signals. Values were normalized as described in Materials and Methods. Obtained ΔOD values are plotted according to age at sampling. Individual animals are distinguished by different colors and longitudinal sera connected by lines. (C) ΔOD values depicted in panels A and B grouped per category. Individual values and mean ± standard deviation (SD) are shown. Statistical significance was tested by Mann-Whitney test using GraphPad Prism: **** indicates p < 0.0001. OD = optical density

### Development of gB and gH/gL ELISAs for EEHV1A, 1B, 4 and 5A

The EEHV1A gB ELISA used to detect EEHV-specific antibody responses in Figure 1 is known to detect antibodies cross-reactive to multiple EEHV (sub)species (3, 5), yet its exact specificity is unknown. Consequently, based on the current data, it is not possible to distinguish whether EEHV-HD fatalities with high gB-specific antibody levels died due to (I) reinfection with or reactivation of the EEHV (sub)species with which the animal had already been infected or (II) primary infection with a heterologous EEHV (sub)species.

To address this question and to assess specificity of our (EEHV1A) gB ELISA, we set out to develop gB and gH/gL ELISAs for all EEHV (sub)species infecting Asian elephants. Both gB and gH/gL are major targets of the EEHV-specific antibody response (3), yet while the gB protein is relatively conserved (64.1 – 95.8% amino acid identity between the different EEHV (sub)species; Table 1 and Figure 2A), the gH/gL dimer is more variable between the EEHV (sub)species (42.0 – 87.0% amino acid identity; Table 2 and Figure 2B). Due to the relatively low level of conservation of gH/gL as compared to gB it is expected to induce lower levels of cross-reactive antibodies upon infection (19) and thereby allow better distinction between serological responses elicited against different EEHV (sub)species.

**Figure 2.**
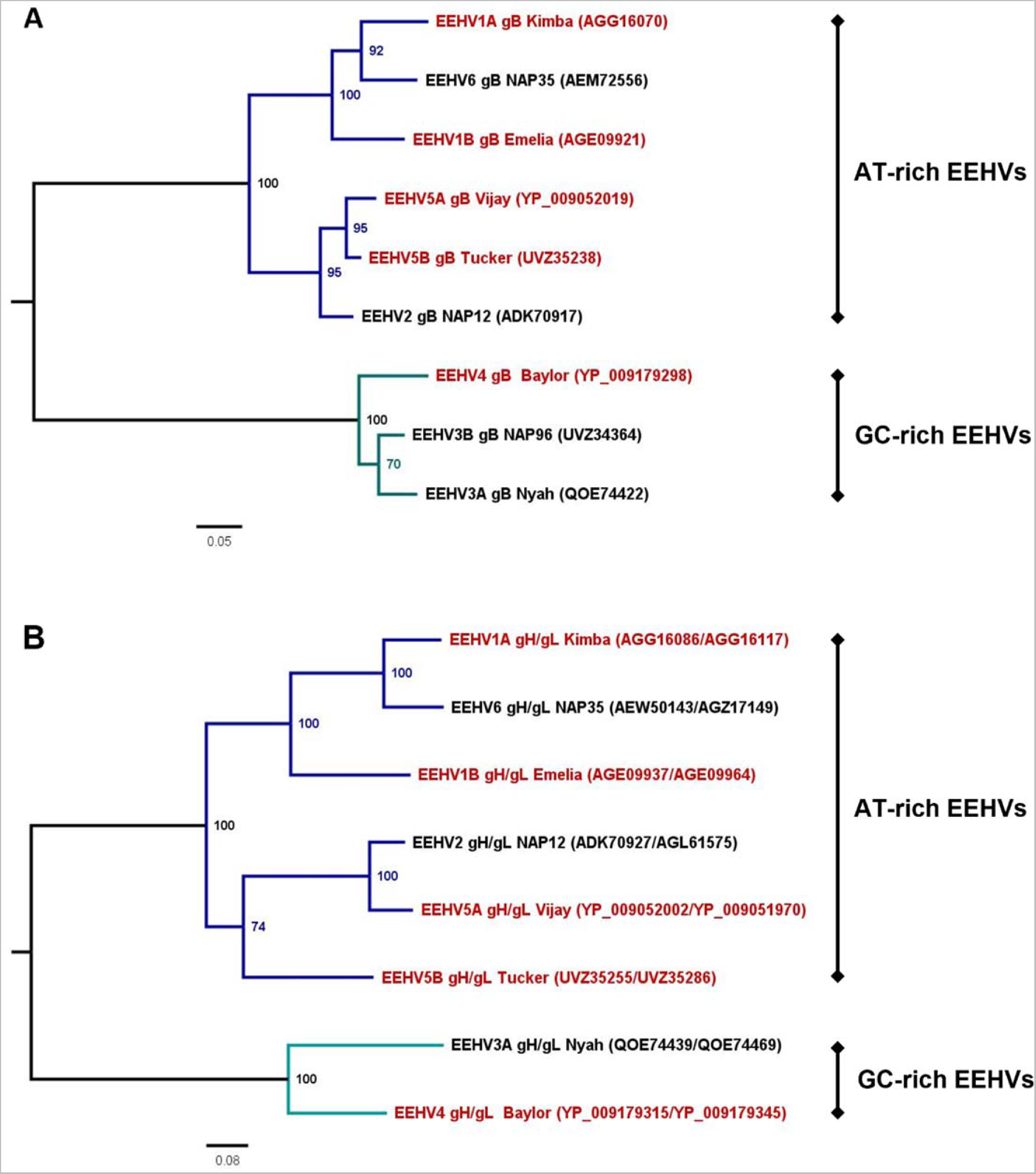
Maximum Likelihood Trees inferred from gB and gH/gL amino acid sequences of different EEHV (sub)species. Full length amino acid sequences of gB (A) and gH and gL (B) were retrieved from GenBank for one viral strain per EEHV subspecies. To facilitate phylogenetic analysis, gH and gL sequences were concatenated. Sequences were aligned using Clustal Omega and trees were constructed using IQTree based the JTT+G4 (gB (A)) and WAG+F+I+G4 (gH/gL (B)) evolutionary model and using 1000 bootstrap replicates. Inferred trees were visualized and edited in FigTree. Presented trees are midpoint rooted. Only bootstrap values ≥ 70 are shown. EEHV (sub)species, viral strain analyzed, and GenBank accession numbers are indicated. EEHV subspecies infecting Asian elephants are highlighted in red; subspecies infecting African elephants are shown in black. Branches including the AT-rich EEHVs are colored dark blue; branches including the GC-rich EEHVs are shown in teal.

**Table 2.**
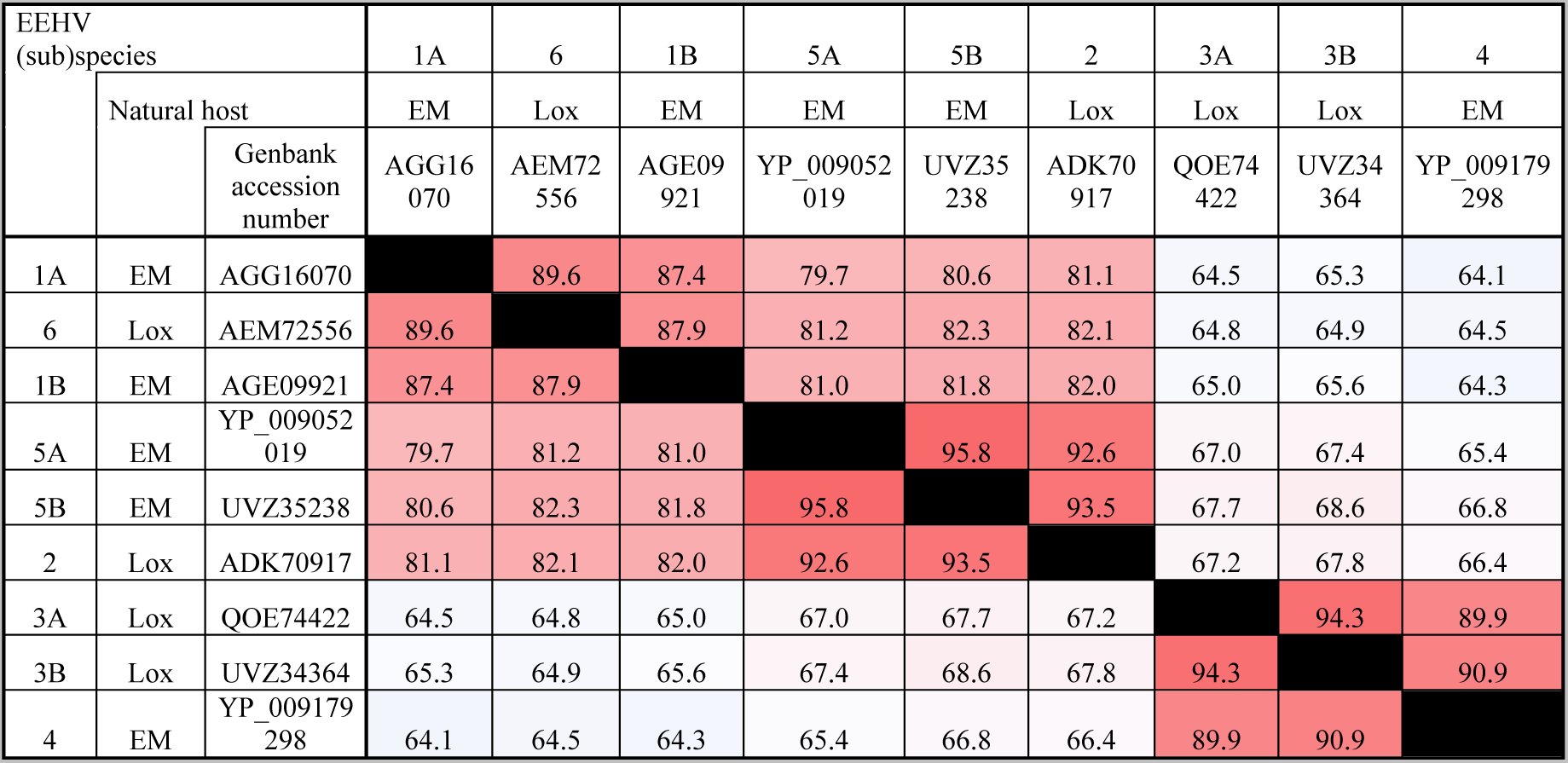
Identity matrix showing gB protein conservation between different EEHV (sub)species. Amino acid identity (in %) is colored based on conservation level, with high conservation shown in red and low conservation shown in blue and the same color scheme being used in Table 2 and 3. EEHV (sub)species, natural host of the (sub)species, and viral strain analyzed are indicated. EM (*Elephas maximus*) refers to the Asian elephant. Lox (*Loxodonta species*) refers to African elephants.

| EEHV (sub)species |  |  | 1A | 6 | 1B | 5A | 5B | 2 | 3A | 3B | 4 |
| --- | --- | --- | --- | --- | --- | --- | --- | --- | --- | --- | --- |
|  | Natural host |  | EM | Lox | EM | EM | EM | Lox | Lox | Lox | EM |
|  |  | Genbank accession number | AGG16070 | AEM72556 | AGE09921 | YP_009052019 | UVZ35238 | ADK70917 | QOE74422 | UVZ34364 | YP_009179298 |
| 1A | EM | AGG16070 |  | 89.6 | 87.4 | 79.7 | 80.6 | 81.1 | 64.5 | 65.3 | 64.1 |
| 6 | Lox | AEM72556 | 89.6 |  | 87.9 | 81.2 | 82.3 | 82.1 | 64.8 | 64.9 | 64.5 |
| 1B | EM | AGE09921 | 87.4 | 87.9 |  | 81.0 | 81.8 | 82.0 | 65.0 | 65.6 | 64.3 |
| 5A | EM | YP_009052019 | 79.7 | 81.2 | 81.0 |  | 95.8 | 92.6 | 67.0 | 67.4 | 65.4 |
| 5B | EM | UVZ35238 | 80.6 | 82.3 | 81.8 | 95.8 |  | 93.5 | 67.7 | 68.6 | 66.8 |
| 2 | Lox | ADK70917 | 81.1 | 82.1 | 82.0 | 92.6 | 93.5 |  | 67.2 | 67.8 | 66.4 |
| 3A | Lox | QOE74422 | 64.5 | 64.8 | 65.0 | 67.0 | 67.7 | 67.2 |  | 94.3 | 89.9 |
| 3B | Lox | UVZ34364 | 65.3 | 64.9 | 65.6 | 67.4 | 68.6 | 67.8 | 94.3 |  | 90.9 |
| 4 | EM | YP_009179298 | 64.1 | 64.5 | 64.3 | 65.4 | 66.8 | 66.4 | 89.9 | 90.9 |  |

DNA constructs for production of soluble recombinant gB, gH and gL of EEHV (sub)species 1B, 4 and 5A were designed according to procedures for EEHV1A (3) and are listed in Table 1. The gB, gH and gL proteins of EEHV5B could not be included since the genetic sequences required became available only recently. Gelcode Blue stained gels containing affinity-purified gB-3×ST and gH-3×ST/gL-6×His proteins (purified based on the StrepTag) are shown in Figure 3A respectively 3B. To improve visualization, gH/gL heterodimer-containing preparations were treated with PNGase F prior to electrophoresis. Clear protein bands at expected molecular weights are visible for all proteins with only minor contaminations being present. Co-purification of gL-6×His with gH-3×ST indicated formation of stable gH/gL heterodimers for all EEHV (sub)species (Figure 3B). In addition, presence of both proteins in fractions affinity purified using the StrepTag was confirmed by detection of their respective protein Tags using Western Blot (Figure 3C).

**Figure 3.**
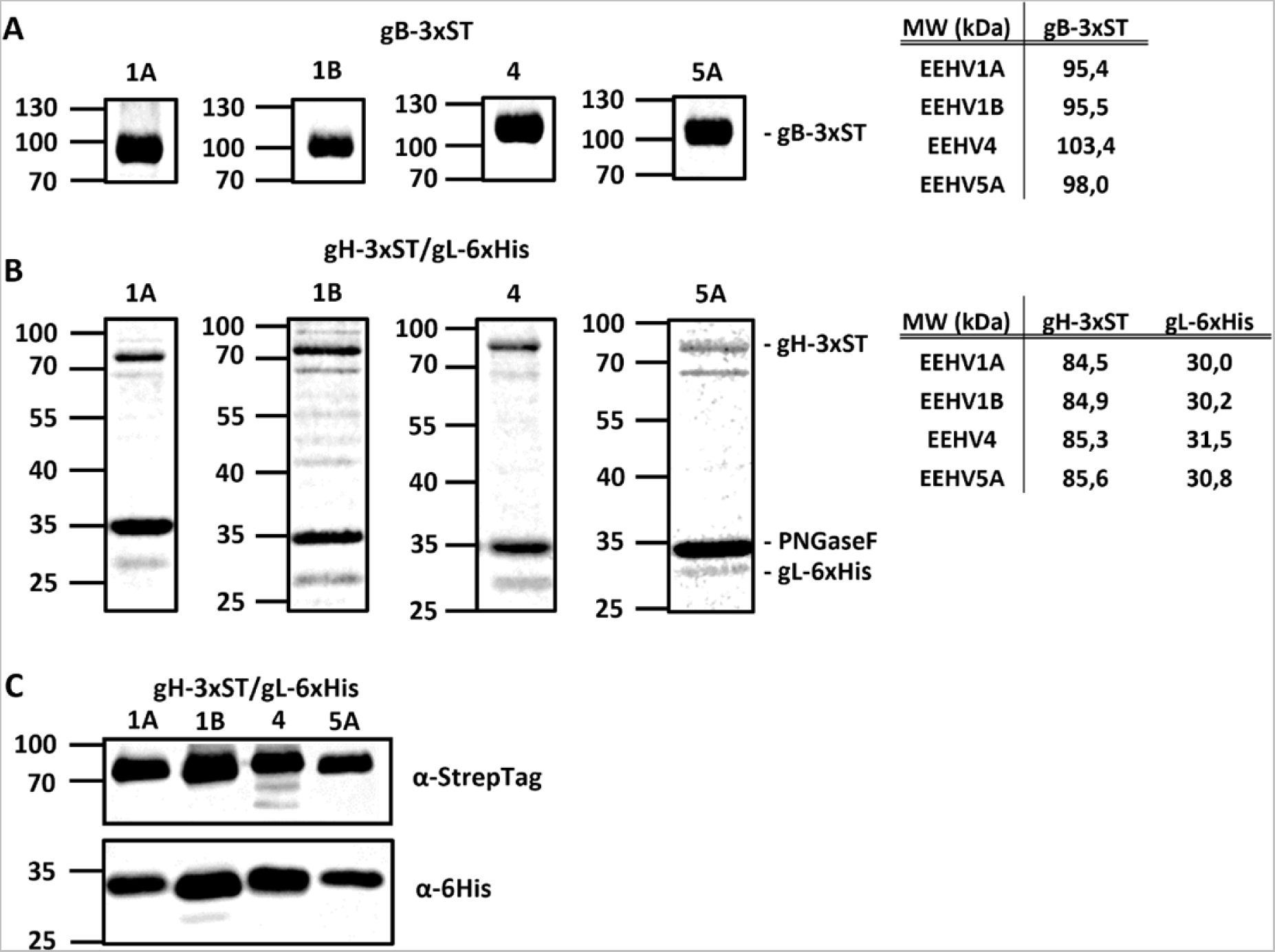
Production of recombinant gB and gH/gL proteins of the Asian elephant EEHV (sub)species. Gelcode Blue-stained gels electrophoresed with gB-3xST (A) and gH-3xST/gL-6xHis (B) for EEHV subspecies 1A, 1B, 4 and 5A affinity purified using the StrepTag. Molecular mass markers are indicated on the left side of the gels. (A) Expected molecular weights of glycosylated gB proteins are listed indicated in the table on the right. (B) gH/gL protein fractions were deglycosylated by PNGaseF (~35kDa) prior to electrophoresis. Expected molecular weights of deglycosylated gH and gL proteins are listed indicated in the table on the right. (C) Western Blot of gels electrophoresed with secreted gH-3xST/gL-6xHis proteins for EEHV subspecies 1A, 1B, 4 and 5A, stained using anti-StrepTag and anti-HisTag antibodies. Protein fractions were deglycosylated by PNGaseF prior to electrophoresis. Expected molecular weights as in panel B.

ELISAs were developed using all antigens according to procedures described previously (3) and subsequently assessed using the subset of sera of elephants <10 years of age included in our cohort at the start of the study (76 sera from 51 animals). In each individual ELISA sera showing clear antigen-specific antibody responses as well as sera without detectable antibody responses could be identified (Figure 4). Mean antibody levels detected for gB of subspecies 1A, 1B and 5A were largely comparable (normalized ΔOD values between 0.57 – 0.66), while the mean antibody level detected for gB of EEHV4 was somewhat lower (normalized ΔOD of 0.32; Figure 4A). Levels of gH/gL-directed antibodies were largely comparable between (sub)species 1A, 4 and 5A (normalized ΔOD values between 0.37 – 0.48), while antibody levels measured using the EEHV1B gH/gL ELISA were lower (normalized ΔOD of 0.15; Figure 4B).

**Figure 4.**
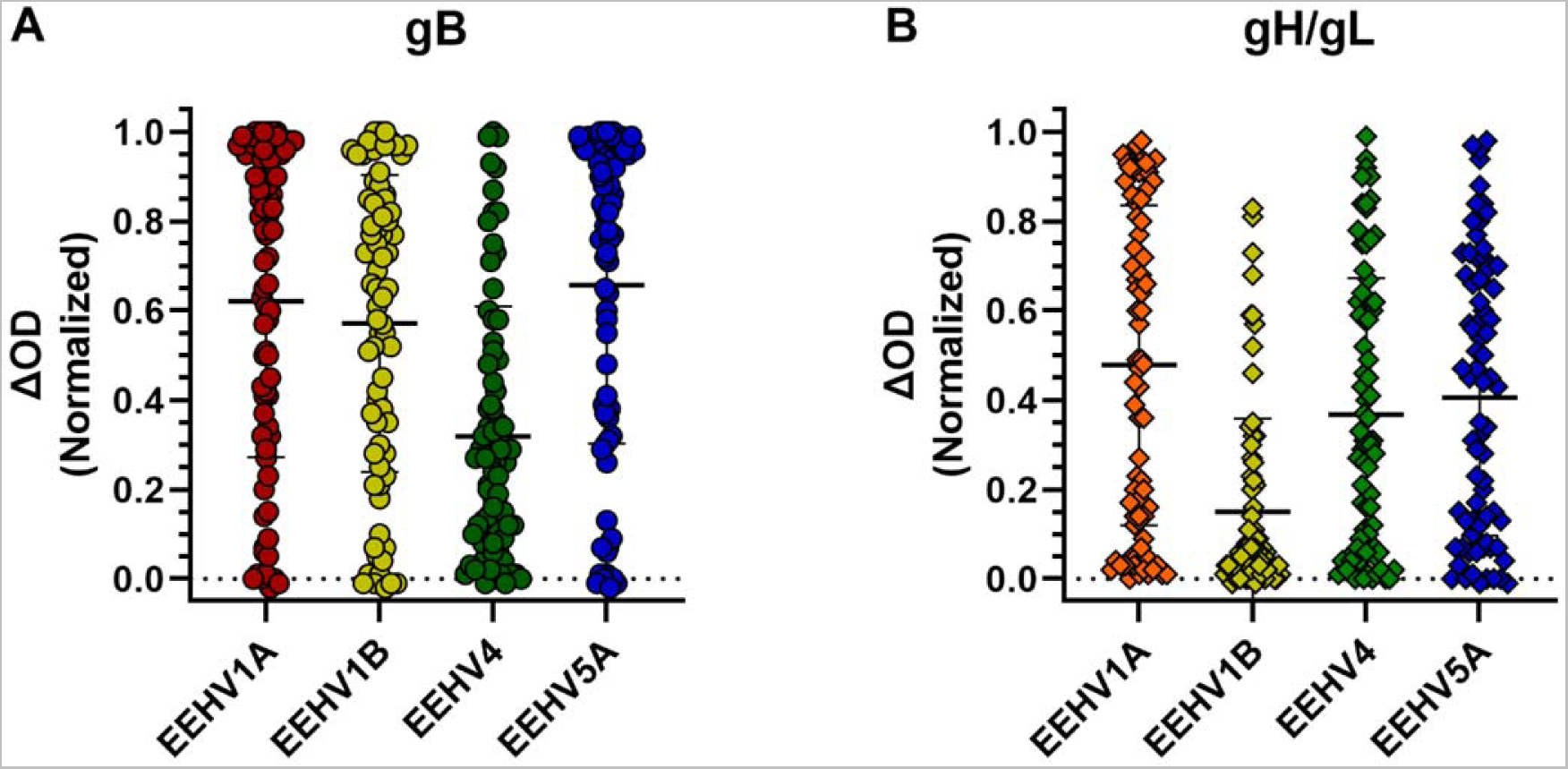
Antibody levels against gB and gH/gL of different EEHV (sub)species measured in sera of Asian elephants < 10 years of age. Antibody levels were measured against gB (A) and gH/gL (B) of all EEHV subspecies for 76 sera of 51 individual animals <10 years of age. A maximum of two sera per animal were included. Samples were tested at a 1:100 dilution and ΔOD values were calculated and normalized as described in materials and methods. Values are grouped per EEHV (sub)species showing both individual values and mean ± standard deviation (SD).

### Correlation amongst gB respectively gH/gL-specific antibody levels of the different EEHV (sub)species

To assess the specificities of the gB and gH/gL ELISAs, it would be preferable to use antisera of animals confirmed positive for one EEHV (sub)species, but negative for all other (sub)species, however these type of predefined sera are not available. As alternative, we assessed to what extent antibody levels measured in ELISAs with antigens of different (sub)species correlated. High correlations can be explained by co-infections and/or presence of cross-reactive antibodies (19). Strong positive correlations were observed between antibody levels detected against gB of EEHV1A, 1B and 5A (Figure 5A, upper panels; R^2^ ≥ 0.89) while only moderate correlations were detected when comparing antibody levels of these subspecies with antibody levels against gB of EEHV4 (Figure 5A, lower panels; R^2^ ≤ 0.43). Notably, the fact that antibody correlation levels were strongly associated with the level of gB conservation between the (sub)species (Figure 5B) indicates that high correlations between antibody levels can largely be explained by antibody cross-reactivity (19).

**Figure 5.**
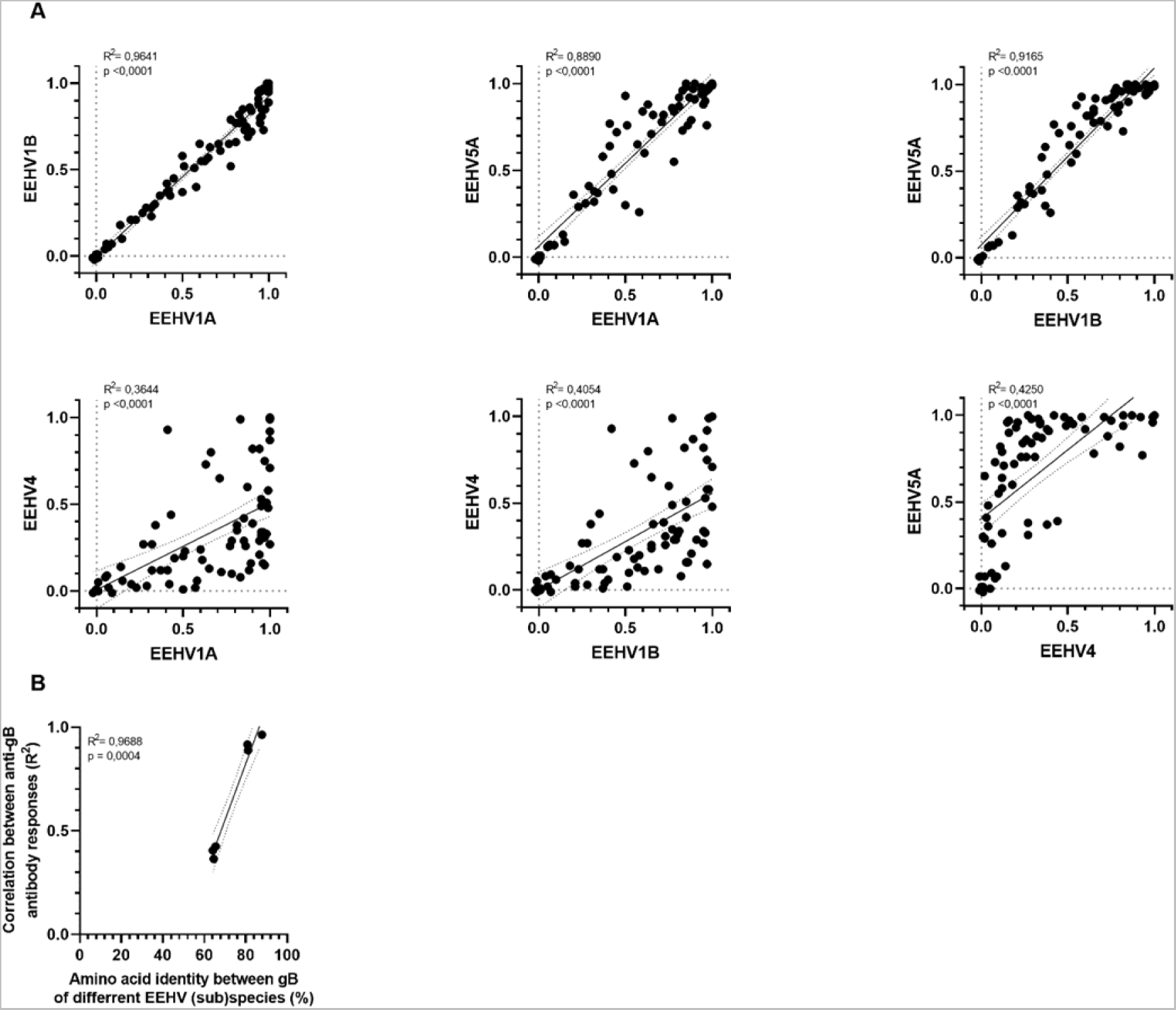
Linear regression analyses of antibody levels against gB of the different EEHV subspecies. (A) Pairwise simple linear regression analyses for ΔOD values presented in Fig. 4A. R^2^ and p-values calculated for each regression are shown in the top left corner of each panel. (B) Correlation between the R^2^ levels obtained in (A) and amino acid identity (in %) as shown in Table 2. Calculated R^2^ and p-values are shown in the top left corner.

Results of pairwise linear regression analyses of observed antibody levels specific for gH/gL of the different EEHV (sub)species are shown in Figure 6A. The (sub)species pairs for which antibody levels against gB were strongly correlated (EEHV1A, EEHV1B and EEHV5A; R^2^ ≥ 0.89), did not show strong correlations between antibody levels against gH/gL (R^2^ = 0.15 – 0.55). Correlations between antibody levels against gH/gL of these (sub)species and EEHV4 (R^2^ = 0.29 – 0.58) were comparable to those observed for gB (R^2^ = 0.36 – 0.42). No relation was observed between correlation levels of gH/gL-directed antibody levels and conservation of gH/gL (concatenated; Figure 6B), or gH or gL solely (Figure 6C respectively 6D). The much lower conservation of gH/gL compared to gB, combined with the general low correlations between antibody responses to gH/gL of different (sub)species, and the absence of a correlation thereof with conservation of gH/gL (in contrast to gB) indicates much less cross-reactive antibodies are elicited against gH/gL than against gB.

**Figure 6.**
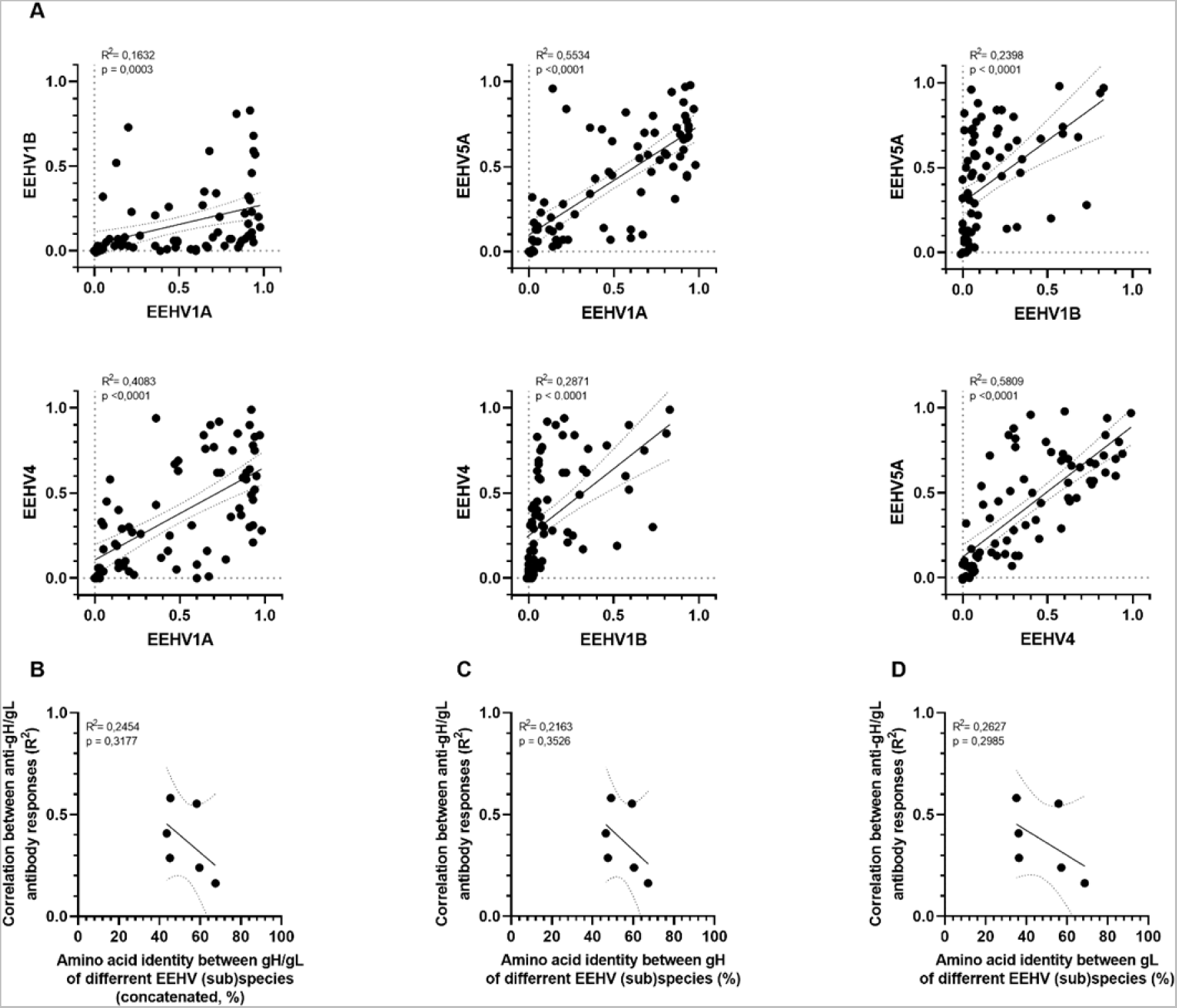
Linear regression analyses of antibody levels against gH/gL of different EEHV subspecies. (A) Pairwise simple linear regression analyses for ΔOD values presented in Fig. 4B. R^2^ and p-values calculated for each regression are shown in the top left corner of each panel. (B-D) Correlation between the R^2^ levels obtained in (A) and pairwise amino acid identity of gH/gL dimer (B; shown in Table 3), gH (C) and gL (D). Calculated R^2^ and p-values are shown in the top left corner of each panel.

**Table 3.**
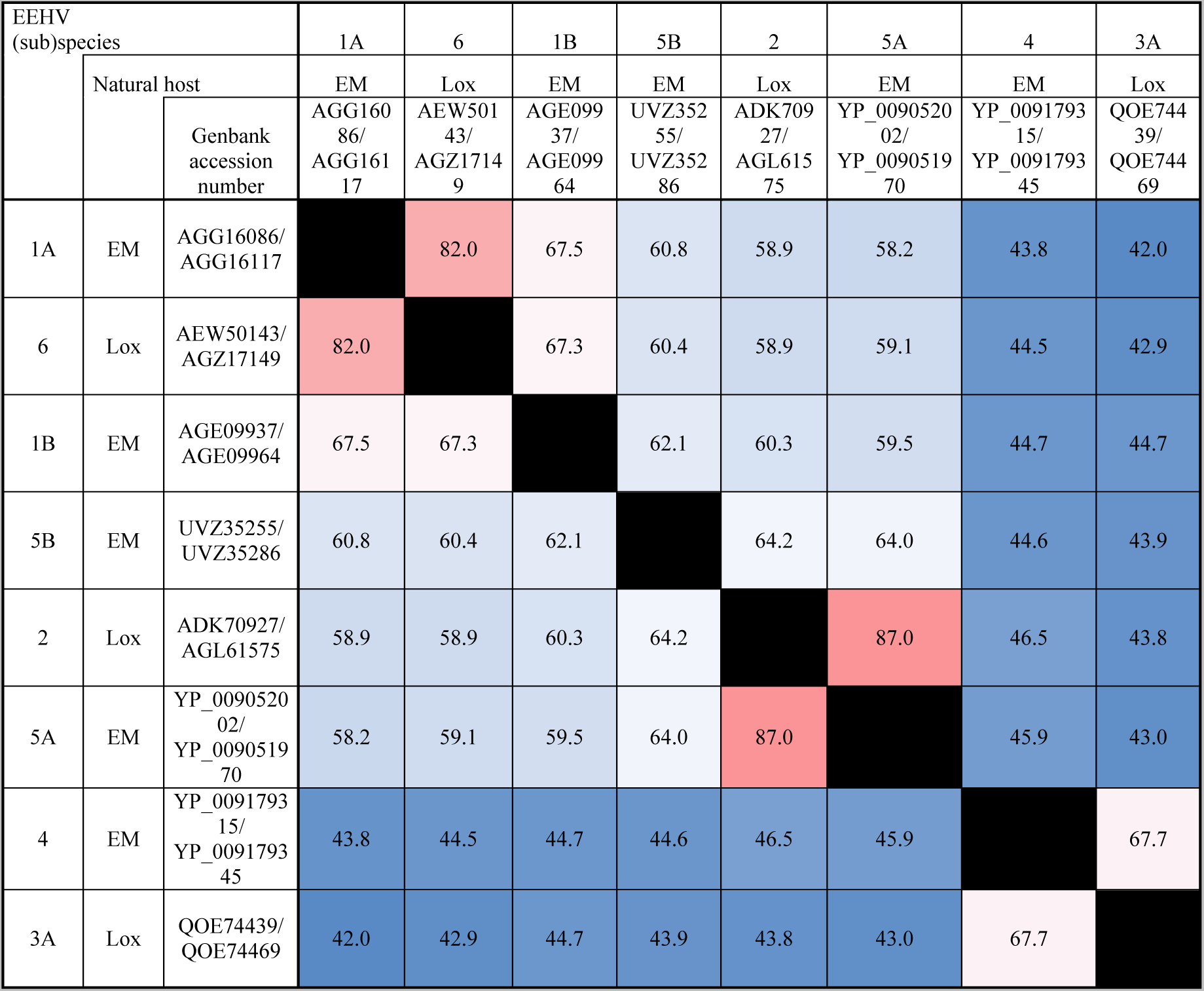
Identity matrix showing gH/gL amino acid conservation between different EEHV subspecies. Full length amino acid sequences of the gH and gL proteins were concatenated per viral strain to facilitate analysis. Data is presented as in Table 2.

| EEHV (sub)species |  |  | 1A | 6 | 1B | 5B | 2 | 5A | 4 | 3A |
| --- | --- | --- | --- | --- | --- | --- | --- | --- | --- | --- |
|  | Natural host |  | EM | Lox | EM | EM | Lox | EM | EM | Lox |
|  | Genbank accession number |  | AGG16086/<br>AGG16117 | AEW50143/<br>AGZ17149 | AGE09937/<br>AGE09964 | UVZ35255/<br>UVZ35286 | ADK70927/<br>AGL61575 | YP_009052002/<br>YP_009051970 | YP_009179315/<br>YP_009179345 | QOE74439/<br>QOE74469 |
| 1A | EM | AGG16086/<br>AGG16117 |  | 82.0 | 67.5 | 60.8 | 58.9 | 58.2 | 43.8 | 42.0 |
| 6 | Lox | AEW50143/<br>AGZ17149 | 82.0 |  | 67.3 | 60.4 | 58.9 | 59.1 | 44.5 | 42.9 |
| 1B | EM | AGE09937/<br>AGE09964 | 67.5 | 67.3 |  | 62.1 | 60.3 | 59.5 | 44.7 | 44.7 |
| 5B | EM | UVZ35255/<br>UVZ35286 | 60.8 | 60.4 | 62.1 |  | 64.2 | 64.0 | 44.6 | 43.9 |
| 2 | Lox | ADK70927/<br>AGL61575 | 58.9 | 58.9 | 60.3 | 64.2 |  | 87.0 | 46.5 | 43.8 |
| 5A | EM | YP_009052002/<br>YP_009051970 | 58.2 | 59.1 | 59.5 | 64.0 | 87.0 |  | 45.9 | 43.0 |
| 4 | EM | YP_009179315/<br>YP_009179345 | 43.8 | 44.5 | 44.7 | 44.6 | 46.5 | 45.9 |  | 67.7 |
| 3A | Lox | QOE74439/<br>QOE74469 | 42.0 | 42.9 | 44.7 | 43.9 | 43.8 | 43.0 | 67.7 |  |

### Antibodies specific for gH/gL of a single EEHV (sub)species were discerned in several animals

To further evaluate whether the gH/gL ELISAs may be used to differentiate between antibodies elicited against different EEHV (sub)species, we assessed gH/gL-specific antibody levels in 199 sera of 80 individual elephants below 10 years of age. In sera of eleven elephants (14%) antibodies primarily directed to one EEHV (sub)species were observed (Figure 7A). For three of these elephants, a prior EEHV infection was confirmed by PCR. In all three cases, the (sub)species against which the most prominent antibody levels in the gH/gL ELISAs was observed corresponded to the (sub)species detected by PCR.

**Figure 7.**
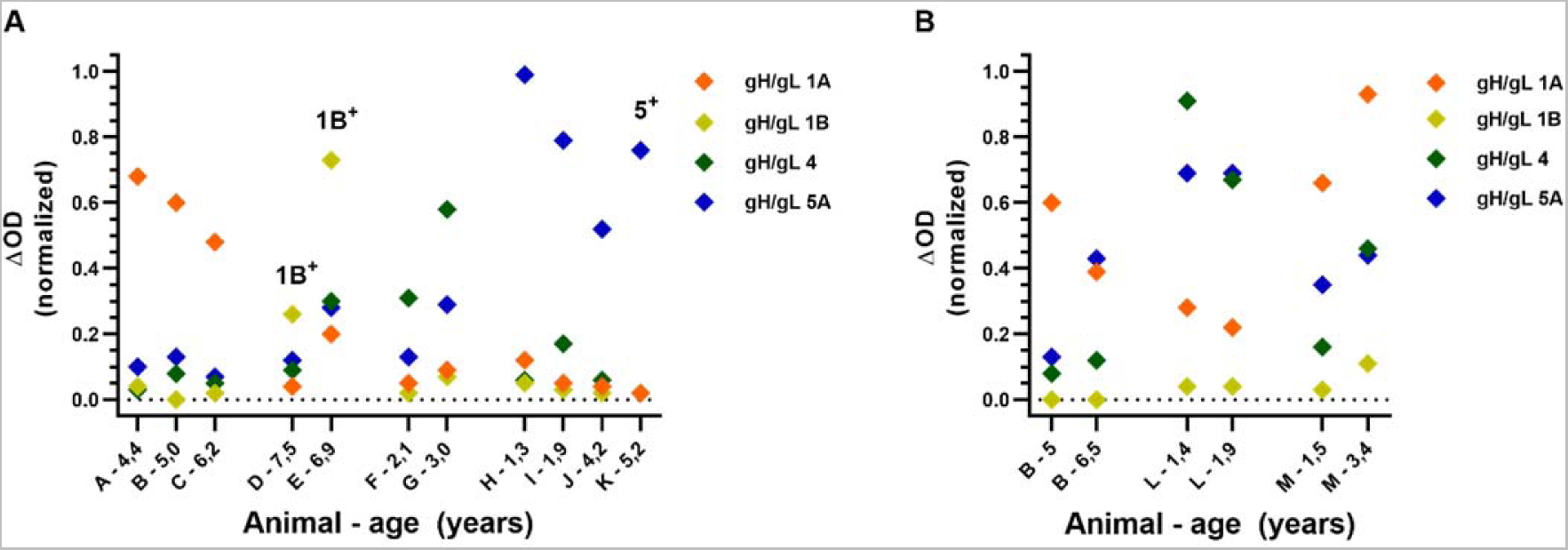
Antibody levels primarily reactive to gH/gL of a particular EEHV (sub)species as detected in sera of Asian elephants < 10 years of age. Normalized ΔOD values detected in sera of 11 individual elephants (A) and paired sera of 3 individual elephants (B) using gH/gL ELISAs for the different EEHV subspecies. Sera of individual elephants are identified by capitals (A-M) and for each serum sample elephant age at time of sampling is indicated. Samples of elephants that previously tested PCR positive for a specific EEHV subspecies are indicated by ((sub)species)+. All samples were tested and ΔOD values were calculated and normalized as described in Fig 4.

If antibody responses are specific to a single EEHV (sub)species no interconnected increase or decrease of antibody levels against different (sub)species would be expected between paired sera. In paired sera of three individual animals (sub)species-specific antibody responses could clearly be discerned (Figure 7B). For animal B, antibody levels against gH/gL of EEHV5A clearly increased during a 1.5-year interval, while antibody levels against EEHV1A decreased and antibody levels against EEHV1B and EEHV4 remained stable. For animal L, antibody levels against gH/gL of EEHV4 decreased during a 5-month interval, while antibody levels against the other (sub)species remained stable. Finally, animal M showed increased antibody levels against gH/gL of all EEHV (sub)species over a 2-year period, yet the extent by which antibody levels increased clearly differed. From these results we conclude that the gH/gL ELISAs are sufficiently specific to distinguish between infections with different EEHV (sub)species, even though we cannot completely exclude that cross-reactive antibody responses against gH/gL of different EEHV (sub)species may (occasionally) be present.

### EEHV-HD fatalities never have high antibody levels to gH/gL of the (sub)species they succumbed to

Next, the gH/gL ELISAs were employed to test sera of 23 EEHV-HD fatalities shown in Figure 1B. The EEHV-HD fatalities for which no gB-specific antibodies were detected in the peri-mortem serum sample, also had virtually non-detectable antibody levels against gH/gL of all (sub)species (n=11; Figure 8A). Moreover, none of the EEHV-HD fatalities positive in the gB ELISA showed high antibody levels against gH/gL of the (sub)species they succumbed to (n=12, Figure 8B-E). Three EEHV1A-HD fatalities for which clear and relatively stable antibody levels to gB were detected, only showed low antibody levels against gH/gL of EEHV1A in all serum samples analyzed (Figure 8B). Likewise, two animals for which virtually no antibodies against gH/gL of EEHV1B were detected, eventually succumbed to EEHV1B-HD (Figure 8C). Figure 8D shows three EEHV-HD cases for which waning of (presumably maternal) EEHV-specific antibodies was detected over time. When these animals succumbed to EEHV1A-HD, antibody levels specific for gH/gL of EEHV1A had waned to low levels (ΔOD ≤ 0.23). Finally, Figure 8E shows antibody levels of four EEHV1A-HD cases for which an increase in EEHV1A gH/gL specific antibodies was observed over time. Even though two animals (case 17 and case 23) had considerable EEHV1A-specific antibody levels in the serum sample taken peri-mortem, all four animals were virtually seronegative to EEHV1A gH/gL in the last serum sample taken before onset of EEHV-HD.

**Figure 8.**
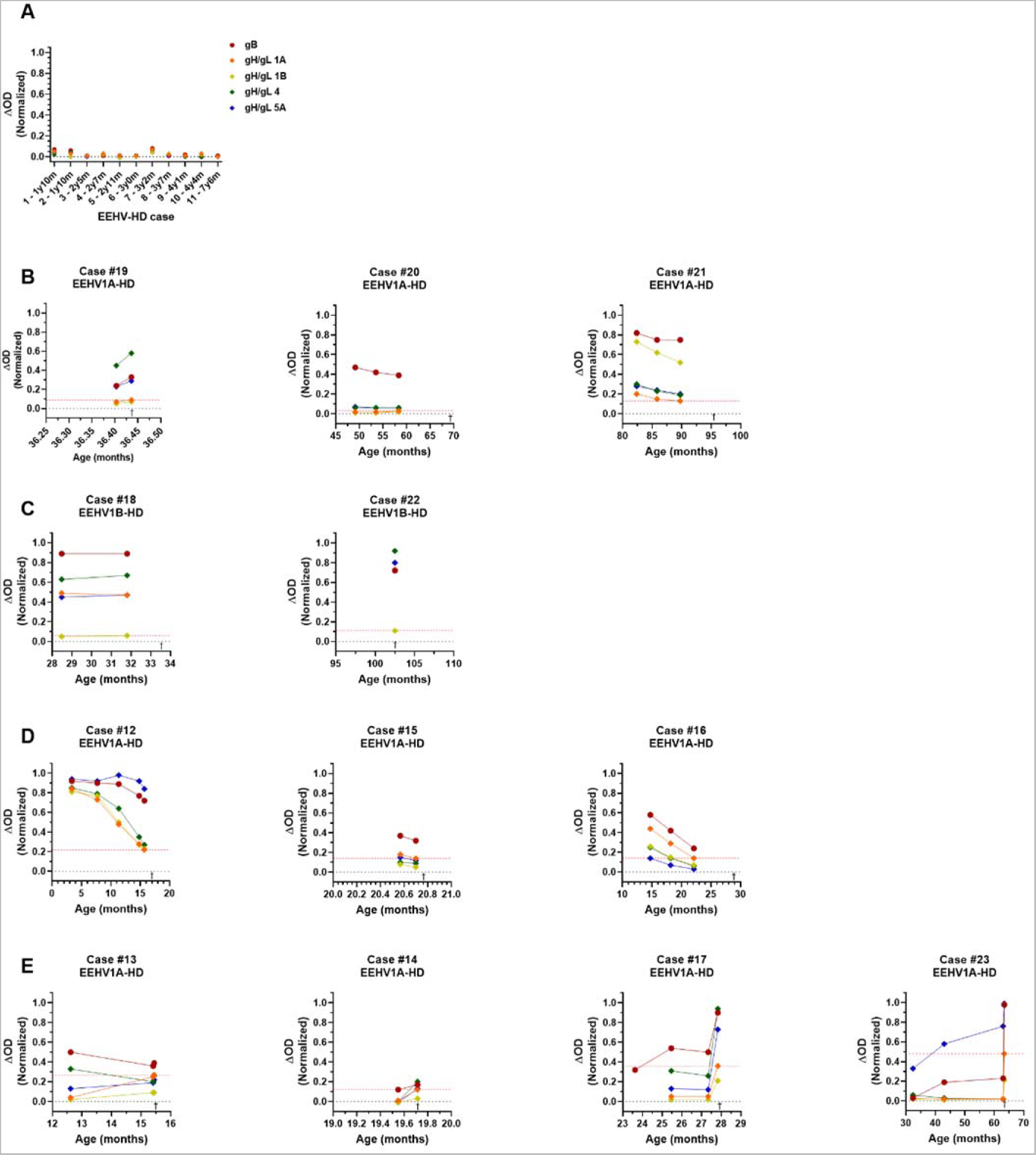
Antibody levels detected in sera of 23 fatal EEHV-HD cases using the multiple EEHV species gB ELISA and (sub)species-specific gH/gL ELISAs. (A) gH/gL specific antibody levels of 11 animals that showed virtually no gB-specific antibody levels in the last serum sample taken before death. EEHV-HD cases shown succumbed to EEHV1A (7 individuals), EEHV1B (2 individuals), or EEHV5 (1 individual). One animal succumbed to a EEHV1A/EEHV4 co-infection. (B-E) Individual panels showing gH/gL-specific antibody levels in (longitudinal) serum samples of 12 EEHV-HD cases with detectable gB-specific antibodies. Respective EEHV-HD cases died due to either a EEHV1A (B, D and E) or EEHV1B (C) infection. In each panel the antibody level against gH/gL of the subspecies the animal succumbed to is indicated by a red dotted line, and time of death is indicated by †. Antibody levels against gB are included for reference purposes. All samples were tested and values were normalized as described previously.

### Seropositivity to gH/gL of the different EEHV (sub)species increases with age

Normalized gH/gL ΔOD values specific for the HD-causing EEHV (sub)species were always below 0.23 in the last sample taken before onset of EEHV-HD (Figure 8). Taking a slightly higher ΔOD value (0.25) as arbitrary cutoff for (putative) protection against EEHV-HD, we next assessed the proportion of animals at risk for developing EEHV-HD per age group (Figure 9A). With exception of calves below 1 year of age, which have pathogen-specific antibody levels comparable to their dams (2, 15), there is an age-related increase in (putative) protection against EEHV-HD caused by the different (sub)species. Between 1 and 5 years of age, when EEHV-HD is primarily observed, a large proportion of animals appears susceptible to EEHV-HD upon infection with most or all (sub)species. In contrast, the vast majority of animals older than 10 seem to be protected against EEHV-HD inflicted by all (sub)species.

**Figure 9.**
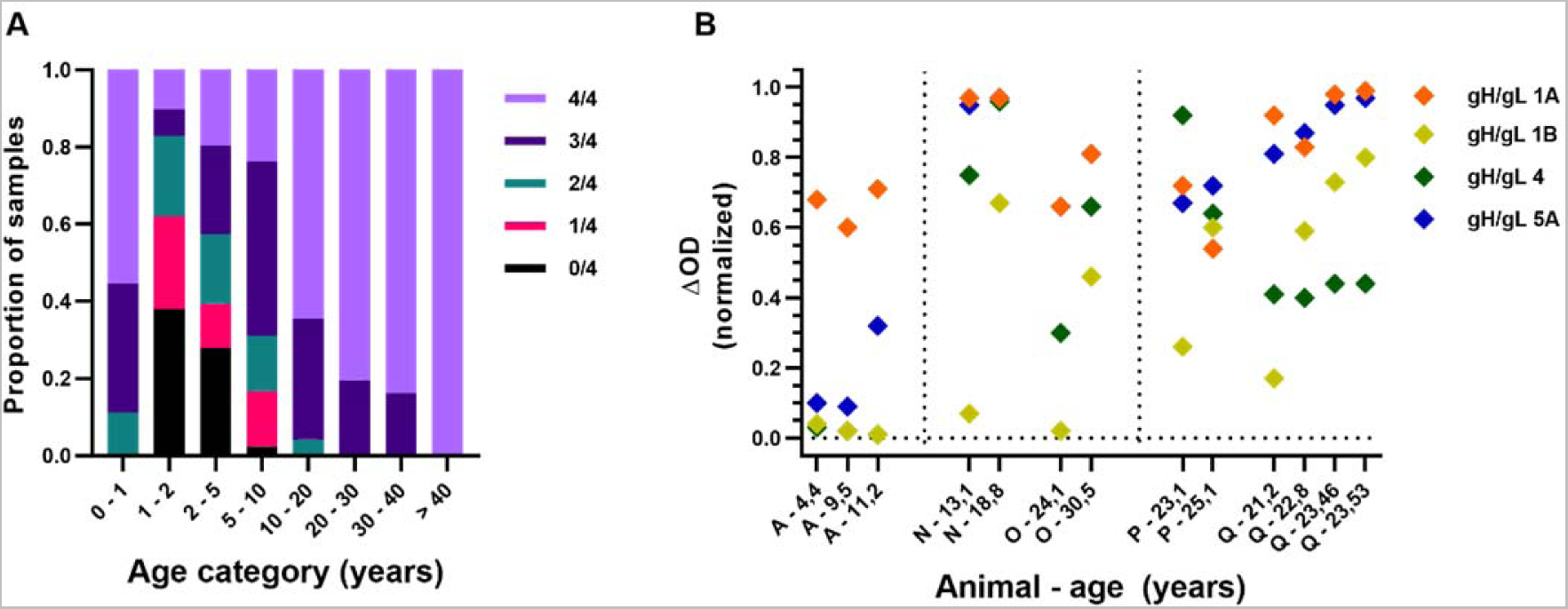
Antibody levels measured against gH/gL of the different EEHV (sub)species with increasing age. (A) Antibody levels against gH/gL of the different EEHV (sub)species was assessed for a total of 298 sera, divided over 8 different age categories. Numbers of samples included per age category ranged from 9 to 61, with no more than two samples per individual animal included per age category. Proportion of samples for which ΔOD levels above an (arbitrary) cutoff level of 0.25 were detected in (n=0-4)/4 gH/gL ELISAs is indicated by different colors. (B) Normalized ΔOD values detected in paired sera of five individual elephants > 10 years of age using gH/gL ELISAs for the different EEHV (sub)species. Individual elephants are identified by a capital letter and for each sample elephant age at time of sampling is indicated. All samples were tested and values were normalized as described previously.

The increased number of EEHV (sub)species to which clear antibody levels (and putative protection against EEHV-HD) is observed may either be caused by infection with various EEHV (sub)species over time and/or by increased levels of cross-reactive antibodies with increasing age, possibly resulting from repeated stimulation of the immune response. We therefore wondered whether (sub)species-specific reactivity could still be observed in older elephants. Figure 9B shows antibody levels of five Asian elephants above 10 years of age for which (sub)species-specific reactivity was indeed observed. Animal A (also shown in Figure 7A) primarily showed reactivity to EEHV1A at both 4.4 and 9.5 years of age. At 11.2 years of age this animal also showed antibodies to EEHV5A, but was still completely unreactive to EEHV1B and EEHV4. Animals N and O, which are part of the same herd and for which the first sample was taken only one month apart, were both (close to) seronegative to EEHV1B at first sampling. In a 5 to 6 year interval, both animals developed antibodies to EEHV1B, suggesting both animals came in contact with this subspecies in the same time period. Reactivity to gH/gL of other EEHV (sub)species also increased, but to different extents. Animal P showed a clear increase in EEHV1B reactivity between 23 and 25 years of age, while reactivity to the other antigens decreased or remained stable. Lastly, animal Q, which is part of the same herd as animal P but which was sampled approximately 15 years later, showed a clear increase of EEHV1B-specific antibodies in a 2-year interval, while reactivity to other antigens was relatively stable. In the same 2-year interval, a fatal EEHV1B-HD case (Figure 8D; case number 8) was observed within this herd, confirming active shedding of EEHV1B within the herd during this period. From this, we conclude that (sub)species-specific serological responses may also be detected in older animals using our gH/gL ELISAs.

## Discussion

In the current study, we developed gB and gH/gL ELISAs for the different EEHV (sub)species infecting Asian elephants and tested their performance using a large panel of elephant sera. Antibody levels measured against gB of different EEHV (sub)species were observed to be strongly correlated, suggesting that antibodies against this antigen are highly cross-reactive between the different EEHV (sub)species. In contrast, antibody levels against gH/gL of the different EEHV (sub)species were far less correlated and hence clearly appear to be more (sub)species-specific. Both gB and gH/gL ELISAs were subsequently used to analyze antibody responses in sera of 23 fatal EEHV-HD cases. Whereas high antibody levels against gB are not necessarily indicative of protection against EEHV-HD, fatal EEHV-HD cases never had high antibody levels against gH/gL of the EEHV (sub)species they succumbed to.

Previous studies already suggested that the gB antigen is recognized by antibodies elicited against multiple EEHV (sub)species (2, 3, 5), yet the exact breadth of gB-specific cross-reactive antibody responses remained unknown. In the current study we observed that antibody levels measured for gB of EEHV1A, 1B and EEHV5A, which all cluster in the AT-rich branch of the EEHV phylogenetic tree (Figure 2A), correlated almost perfectly. Moderate correlations were observed between antibody levels against gB of the AT-rich EEHV (sub)species and EEHV4, which belongs to the GC-rich branch of the EEHV phylogenetic tree. The correlations between antibody levels measured against gB of the different EEHV (sub)species were highly associated with the degree of gB conservation between these (sub)species. Consequently, the strong correlations observed most likely reflect presence of high levels of cross-reactive antibodies, particularly between EEHVs of the AT-rich branch. The extent to which cross-reactive antibodies recognizing gB of both AT-rich and GC-rich EEHV (sub)species are formed cannot be assessed on basis of the current analyses. Yet, this type of antibodies is clearly generated to a much lower extent than gB-specific cross-reactive antibodies between species of the AT-rich branch.

Much lower correlations were observed when comparing antibody responses against gH/gL of different EEHV (sub)species. Moreover, the level of correlation observed could not be explained by protein conservation. Notably, in sera of 11 out of 80 elephants analyzed (14%) antibodies primarily recognizing gH/gL of one EEHV (sub)species were observed. Three of these animals had a prior PCR-confirmed EEHV infection. In all three instances, the (sub)species detected by PCR was in line with the (sub)species against which the most prominent antibody levels were detected using the gH/gL ELISAs. Furthermore, (sub)species-specific antibody kinetics could be observed in both young and adult animals when testing longitudinal serum samples using the novel gH/gL ELISAs.

The data presented above imply that the gH/gL ELISAs have greatly increased specificity for the different EEHV (sub)species over gB ELISAs. However, antibody responses against gH/gL were observed not to be entirely (sub)species-specific. Nearly all animals that predominantly showed antibodies to gH/gL of one (sub)species (Figure 7A), also showed low but detectable antibody levels to gH/gL of at least one other EEHV (sub)species. Additionally, upon acute (and eventually fatal) EEHV1A infection of both case 17 and 23 a clear increase in gH/gL-specific antibody levels was detected for multiple EEHV (sub)species, while to the best of our knowledge no other EEHV (sub)species than 1A was detected during disease. Before onset of EEHV1A-HD, both cases already showed clear antibody levels to gH/gL of other EEHV (sub)species than 1A, indicative of a previous heterologous EEHV infection, and this pre-existing immunity may have contributed to the cross-reactive nature of the observed antibody response. Whether these kind of cross-reactive gH/gL-targeted responses may become more specific with time, for example due to affinity maturation, or remain cross-reactive is currently unknown.

Even though the gH/gL ELISAs are not completely (sub)species-specific, their value was clear when testing 23 fatal EEHV-HD cases. While in 12/23 EEHV-HD fatalities gB-specific antibodies could clearly be detected, none of these fatalities had high antibody levels to gH/gL of the EEHV (sub)species they succumbed to. For at least seven of the EEHV-HD fatalities tested, clearly detectable antibody levels against gH/gL of at least one EEHV (sub)species other than the (sub)species causing EEHV-HD indicate that these animals were already infected with a heterologous EEHV (sub)species before succumbing to EEHV-HD. For example, case 19 (Figure 8B) showed clear antibody levels to both EEHV4 and EEHV5A before succumbing to EEHV1A-HD. Likewise, case 17 (Figure 8E) showed clear antibody levels to EEHV4 and case 23 (Figure 8E) showed clear antibody levels to EEHV5A before they succumbed to EEHV1A-HD. Notably, both EEHV1B fatalities shown in Figure 8C had clear antibody levels against all EEHV (sub)species but EEHV1B, and still succumbed to EEHV1B infection. Overall, these results imply that a previous infection with one EEHV (sub)species does not necessarily protect against EEHV-HD upon infection with another EEHV (sub)species.

Notably, considering the relatively stable antibody levels against gB and the age of the animal at sampling, also case 20 (Figure 8B) had most likely been infected with at least one EEHV (sub)species in the past, yet virtually no gH/gL-specific antibody levels were observed in sera of this animal. It may thus be possible that some animals do not develop clear gH/gL-specific responses upon infection. Alternatively, potentially this animal may have previously been infected with EEHV5B, since this (sub)species could not be included in the current panel of ELISAs.

The chance an animal has antibodies against gH/gL of multiple EEHV (sub)species increases with age. Over 95% of animals above 10 years of age show clear antibody levels to at least 3 EEHV (sub)species, while all animals above 40 are found to have high antibody levels to all EEHV (sub)species. These observations suggest that elephants get infected by virtually all EEHV (sub)species during their lives; a hypothesis supported by the observation that (sub)species-specific gH/gL-directed antibodies may still be observed in animals above 10 years of age. Yet, we cannot completely exclude that repeated stimulation of the immune system may also lead to increased cross-reactivity of gH/gL-directed antibody responses and thereby clearly detectable antibody levels to all EEHV (sub)species. Regardless of the origins of these antibodies, clearly detectable antibodies to virtually all EEHV (sub)species observed in adult animals correlate with the fact that EEHV-HD is almost never observed in elephants over 10 years of age.

Most current efforts to develop an EEHV-HD vaccine make use of gB as the vaccine antigen (20, 21), however gB may not be the most suitable antigen for this purpose. Both in the current and previous studies it was shown that high antibody levels against gB are not necessarily indicative of protection against EEHV-HD (2, 5). Additionally, similar observations were made in clinical trials of vaccine candidates against several human herpesviruses (22, 23). The lack of correlation between gB-specific antibodies and protection may be caused by the (active) prefusion conformation of gB being highly instable and not easily stabilized. As a consequence, most gB molecules will readily adopt the postfusion conformation and the majority of antibodies against gB will thus be formed against this (inactive) postfusion form of gB and are likely not protective. Since antibody responses to gH/gL do correlate with protection against EEHV-HD, we believe gH/gL is a more optimal antigen for EEHV vaccine design.

The data presented in this study strongly suggests that young elephants with low to non-detectable antibody levels to gH/gL of a specific EEHV (sub)species are at risk to develop fatal EEHV-HD upon infection with that particular (sub)species. All 23 EEHV-HD fatalities within the current cohort had antibody levels below (a normalized ΔOD of) 0.23 against gH/gL of the HD-causing EEHV (sub)species just before they developed fatal disease. To the best of our knowledge, a total of 41 Asian elephants succumbed to EEHV-HD in Europe to date, making our cohort of 23 fatalities (56% of the total reported cases) highly representative. The exact antibody level above which animals are protected against EEHV-HD remains to be determined and will be subject to further study.

